# Rapid Carbon Dioxide Capture and Short-Term Biocompatible Sequestration in Aquatic Environments by Monoethanolamine Scrubbing within Calcium Alginate Gel

**DOI:** 10.1101/2022.01.31.478517

**Authors:** Simon Ogundare

**Affiliations:** Department of Chemistry, Columbia University, New York, New York, USA

## Abstract

Alginate is a biopolymer extracted from the cell walls of algae, and can crosslink with divalent cations to form an insoluble hydrogel. In this paper, we develop a method to immobilize monoethanolamine, an amine CO_2_ scrubber, within calcium alginate gel. By mixing monoethanolamine into an alginate solution as the gel was formed, we suspended the compound in the gel, facilitating a means to capture carbon dioxide directly from aquatic environments into the gel, while tethering monoethanolamine and the products formed from CO_2_ capture to the gel. To delay the eventual diffusion of monoethanolamine out of the gel, we investigated (1) the effect of increasing alginate concentration and (2) the effect of additional alginate layers on the outward diffusion of dye placed in the center of the bead. Using ultraviolet-visible spectroscopy to quantify diffusion rates over time, we determined that increased alginate concentration paired with increased layering significantly decreased the rate of outward diffusion. Finally, we prepared beads using North Atlantic seawater as a solvent and compared the rate of dye leakage in seawater and distilled water to that in beads prepared in distilled water. Expectedly, we concluded that beads prepared with solvents isotonic to their environments would exhibit less leakage as well as greater mechanical stability, resisting swelling, bursting, or splitting behaviors.

## Introduction

The anthropogenic release of carbon dioxide is unmistakably a major contributor in the phenomenon of global warming (Inter-governmental Panel on Climate Change, 1990). As a global issue, no one party can be wholly responsible for the implementation of strategies for adaptation or mitigation. While effective technologies currently exist and many are widely known and continually developed, the applications of such methods are often undercut by the accessibility of these methods to the countries, companies, or corporations that can afford and adopt them without significant risk and financial strain. Moreover, as is the case at the intersection of scientific theory and practice, the portrayal and public reception of these geoengineering methods can serve as a social barrier barring greater implementation and is a concern worthy of mention (Westoby & McNamara, 2019).

The aim of this paper is to experimentally combine the effectiveness of monoethanolamine, an amine-based carbon dioxide scrubber, with the biocompatibility and natural accessibility of alginate, a component of macroalgal cell walls, and propose a novel unity of these two systems by developing and optimizing a comparably effective and widely accessible mode of safe, sustainable carbon capture and storage in aquatic environments.

### Monoethanolamine and CO_2_ Scrubbing

Monoethanolamine (MEA), an amine scrubbing agent, has been a popular and prevalent choice for carbon dioxide scrubbing over many years. MEA has been praised for both its fast reaction kinetics with CO_2_, and the relative stability of its generated products. The amine is utilized in industry as an aqueous solvent for the sweetening of flue gas — the scrubbing of gases such as CO_2_ and H_2_S from a homogenous mixture of exhaust gases, including N_2_, NO_x_, O_2_, and H_2_O (Song et al., 2004).

In the process of scrubbing carbon dioxide, two equivalents of MEA react with a molecule of CO_2_. Based on stoichiometry of the reaction, a theoretical maximum of 0.5 moles of CO_2_ can be scrubbed from 1 reacting mole of MEA, as represented by the following reaction:

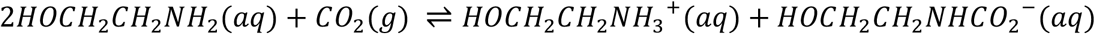

Despite the scrubbing effectiveness of the amine, some crucial limitations stand in the way of its increased use as a carbon dioxide scrubber. Regeneration of MEA and consequent release of stored CO_2_ requires high temperatures, a high energy and high-cost investment for those that intend to release CO_2_ post-capture. Furthermore, in the highly oxidizing environments generated by the chemical constituents of gas streams, amine degradation by oxidation is common – effectiveness can therefore decrease as usage increases (Luis, 2016). Finally, the typical synthesis of the amine requires a reaction between aqueous ammonia and ethylene gas at 50-70 bar (Thiyagarajan et al., 2021), which not only produces MEA, but a ratio of diethanolamine (DEA) and triethanolamine (TEA) yield-dependent on amine concentrations, with both DEA and TEA being relatively inefficient scrubbing compounds, toxic to aquatic organisms, and suspected carcinogens (Libralato et al., 2010). Increase of amine concentrations favors selective production of MEA; however, MEA is not exempt from exposure hazards as the prior article reports the amine as acutely toxic, discouraging the naked release of the compound directly into aquatic environments.

These presented limitations are issues that can be solved once we reframe the objectives of MEA scrubbing — through modifying the very medium in which scrubbing occurs. In this paper, we reframe MEA scrubbing of CO_2_ within the physical scaffolding of the alginate biopolymer, effectively functionalizing the biocompatible gel for carbon capture in aquatic environments.

### Substrate Encapsulation through Alginate

Alginate is extracted from macroalgal species, notable for their rapid rate of growth. Alginate extraction occurs by converting insoluble alginate salts present in the cell walls of algae into water-soluble salts, namely sodium alginate by treating with a base such as NaCO_3_. The extracted sodium salt is then precipitated and worked up to purity (Peteiro, 2018).

Alginate (Alg) is a polysaccharide consisting of two main residues, α-d-mannuronic acid (M) and β-l-guluronic acid (G) connected by 1-4 glycosidic linkages. The constituent residues of the polymer vary in sequence within the alginate polymer, and are seen in groups of M-blocks, with consecutive M subunits, G-blocks, with consecutive G subunits, or MG blocks, with alternating M-G subunits. The M/G ratio has been widely explored in literature, and it has been frequently suggested that an increased proportion of G blocks within a sample of alginic acid results in greater crosslinking and gelation with divalent cations (Peteiro, 2018). However, alginates containing a high M/G ratio have been demonstrated to be more flexible, suitable for heavy metal adsorption, offering a practical application for alginates of this composition (Wang et al., 2016).

Protonated calcium alginate has been recognized by prior literature for its ability to adsorb heavy metals in exchange for protons, and can be regarded as an excellent means for heavy metal scrubbing (Ibáñez & Umetsu, 2002). While heavy metal removal is not significantly explored in this paper, there are noted advantages of the use of calcium alginate in aquatic environments and ecosystems where heavy metal poisoning poses a toxic risk (Papageorgiou et al., 2006).

Sodium alginate is the water-soluble sodium salt of alginic acid. Introducing a source of divalent calcium ions such as CaCl_2_ results in crosslinking as the calcium ions ionically associate with the Alg blocks in an arrangement following the egg-box model (Cao et al., 2020), forming the insoluble gel calcium alginate (Fig. 1). Calcium is commonly used in literature and forms the calcium alginate hydrogel; however, a variety of divalent species, such as barium, can also be used in crosslinking.

**Figure 1:**
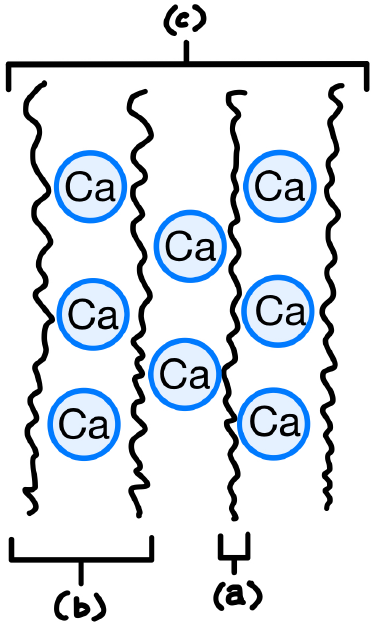
A diagram of the egg-box model used to describe the gelation of calcium alginate gel. Label (a) represents the alginate biopolymer, (b) represents a calcium alginate dimer, with three divalent calcium ions packed against two alginate strands, and (c) represents the multimer.

The physical properties of alginate are commonly examined in the context of the insoluble hydrogel, through gelation. Sodium alginate dispensed in a source of divalent cations (most common among these are Ca^2+^ ions) results in formation of the calcium alginate hydrogel. In CaCl_2_, with a high degree of dissociation, gelation occurs rapidly as calcium displaces sodium in solution and an insoluble hydrogel, calcium alginate, is formed. In solutions that are minimally soluble in water, such as calcium lactate, the rate of gelation decreases as Ca^2+^ is minimally available and released gradually from the lactate.

Many of the promising applications of alginate in its unmodified form ranges from commercial food science to the formation of matrices for encapsulation of enzymes and cells. However, prior literature also explores the potential for chemical modification of alginate, based upon the availability of free functional groups: hydroxyl and carboxyl groups present on the alginate polymer (Yang et al., 2011).

This paper details the preparation of calcium alginate hydrogel, functionalized for CO_2_ scrubbing. In the study, we:

1. Form the alginate hydrogel containing a mixture of sodium alginate with monoethanolamine (MEA) and isolate its native capacity for CO_2_ absorption.
2. Use repeat gelation and increased alginate concentration to significantly delay the leakage of dye out from the core layer of beads.
3. Prepare the alginate gel using North Atlantic Ocean seawater as a solvent to investigate mechanical stability in aquatic environments and assess necessity of solvent distillation.

## Methods

### Experimental Aims

This study was designed to primarily determine the native capacity of monoethanolamine (MEA) for CO_2_ absorption when entrapped in calcium alginate gel, while investigating the impact of multilayering and altering alginate concentration to significantly delay the outward diffusion of MEA through the hydrogel. Finally, to investigate the practicality of the method in the field, a sample of North Atlantic seawater was taken to prepare the alginate hydrogel, and its mechanical stability was compared in saltwater and distilled water through ultraviolet-visible spectrophotometry and compared to a control.

### Accumulation of Carbon Dioxide by Mass in MEA-Alginate Beads

Analytical-grade purities of monoethanolamine (MEA) and sodium alginate were obtained, and reagents were used without any further purification.

A 1:1 volumetric mixture of 30% w/v MEA and 4% w/v sodium alginate were combined using a magnetic stirrer, diluting the concentrations by half (15% and 2%, respectively). The mixture was covered with Parafilm to minimize loss of MEA due to vaporization. The mixture was left for 24 hours to settle and remove stray air bubbles. 25 mL of the mixture was dispensed dropwise via a needleless syringe into 50 mL of a 1.5% w/v calcium chloride solution which was slowly stirred to ensure an even distribution of crosslinking. Once the mixture had been dispensed and the beads were formed, the beads were lightly washed to remove traces of free calcium ions.

The beads were transferred to a vacuum-insulated metal flask, containing 200 mL of distilled water. A phone with a wireless microphone function was placed with the flask on a mass balance, and the total mass of the phone and flask was tared. Masses of dry ice (n = 6) were added to the flask, and the mass displayed on the balance as the dry ice sublimated was recorded every 10 seconds. As the bubbling through the solution emitted sound conducted through the metal of the insulated flask and mass balance, the wireless microphone function was used to hear sublimation and detect when the dry ice stopped sublimating through the solution. At this endpoint, data was collected for another 120 seconds to ensure complete sublimation before adding another mass of dry ice. Minimal condensation appeared at the top of the flask, and this moisture was wiped between runs to ensure the accumulation of mass was restricted to the net gain of dissolved CO_2_ in the flask, rather than from excess moisture from the laboratory.

This experiment was carried out in triplicate, and results were compared to a blank containing 200 mL of distilled water. This was to account for any possible background change in mass by carbon dioxide dissolution into the surrounding water, and to ensure that differences between the blank and the MEA-alginate bead solution were restricted to absorption by MEA. The endpoints were also compared to a blank containing a 0.3 M solution of calcium chloride, to account for background CO_2_ absorption as CaCO_3_ synthesis by free Ca^2+^ ions.

### Effect of Multilayering on Delaying Outward Diffusion of Methylene Blue Dye

One mixture of 0.2% w/v methylene blue dye and 2% sodium alginate and another mixture of 0.2% methylene blue and 4% sodium alginate were combined using magnetic stir bars and left for 24 hours to settle. The mixtures were each dispensed dropwise into separate solutions of 0.3 M calcium chloride under a magnetic stirrer. Of the calcium alginate beads formed from the 2% SA solution, half (n = 3) were randomly selected and placed in three 20 mL scintillation vials, while the other half (n = 3) was placed back in the 2% sodium alginate solution for additional layering. The multilayer beads were transferred from the alginate into a 0.3 M calcium chloride solution, forming a second layer, and this gelation process was repeated to form a third crosslinked layer. The same process was repeated with the mixture containing 4% sodium alginate. In sum, monolayer calcium alginate beads and trilayer calcium alginate beads were formed at w/v concentrations of 2% and 4%.

Once all beads had been transferred to vials and the vials were filled with 7 mL of distilled water, the solution surrounding each bead was transferred to a 1 cm cuvette and measured spectrophotometrically using a Vernier LabQuest 2 and Vernier SpectroVis spectrophotometer.

Spectrophotometer measurements were taken every 30 minutes. The spectrophotometer was re-calibrated with a blank cuvette every half hour before each set of measurements to mitigate systematic deviations across the 8-hour period of measurements, and the exterior of each cuvette used was rinsed with distilled water and wiped with Kim-wipes. Each cuvette was used exclusively for one vial, to prevent mixing of solutions in the cuvettes and unintended dilution or concentration of methylene blue.

This experiment was carried out in triplicate. Results obtained were compared to a blank containing distilled water to establish baseline absorbance values. Results from the 2% alginate beads were compared to a solution containing a drop of the 2% sodium alginate and 0.2% methylene blue, left for 24 hours to diffuse, to establish a maximum absorbance value. The same was repeated for the 4% alginate beads, using drops of the 4% sodium alginate-0.2% methylene blue solution.

### Effect of Aquatic Environments on Mechanical Stability

To investigate the stability of calcium alginate beads in different aquatic environments, a sample of Atlantic Subarctic Upper Water (ASUW), with pH 7.68, salinity of 30 ppt, and specific gravity of 1.023 (within the ASUW range) was used as a solvent to prepare 2% trilayer beads (saltwater beads). These beads were compared to 2% trilayer beads prepared in distilled water solvent, as a proxy for freshwater conditions. The saltwater beads were placed in scintillation vials containing either 7 ml of ASUW or distilled water, and distilled water beads were placed in scintillation vials containing either 7 mL of ASUW or distilled water.

The procedure for taking spectrophotometer measurements was repeated in triplicate. Maximum absorbance values were obtained by dispensing a drop of the saltwater or distilled water solution into a vial containing 7 ml of ASUW or distilled water, respectively. Solutions were agitated so the dye could fully disperse, and absorbance was measured after 5 minutes of agitation. Over an eight-hour period, measurements were reported every 30 minutes. For clarity, graphs report measurements every hour.

### Data and Analysis

For measurements of accumulation of mass by gain of CO_2_, a mass balance was used, and data was recorded by hand every 10 seconds, taking a total of 6070 seconds per trial. The data collected allowed for comparisons of net mass accumulation after complete dry ice sublimation, as well as instantaneous rates of sublimation. Comparisons between total mass of dry ice added across blank and flask containing MEA-alginate beads were also established. Finally, comparisons between total sublimation times for the blank and the flask containing the MEA-alginate beads were established.

Every mass of dry ice dispensed into the flask was cut cylindrically with a diameter of 1.5 centimeters.

Spectrophotometric measurements were recorded on the Vernier Graphical Analysis application, and data was streamed wirelessly from the sensor. Full spectra were reported for reference, yet only the absorption at λ_max_ for methylene blue (667 nm) was used as an indicator of concentration as the dye diffused from the core of the bead to the surrounding solution based on prior literature (Morgounova et al., 2013). The Beer-Lambert law was used to determine the extinction coefficient and ultimately the concentration of the dye at these time intervals, which was used to determine the rates of dye leakage from the monolayer and trilayer calcium alginate beads.

A Mettler Toledo FiveEasy pH meter was used to estimate pH of the Atlantic Subarctic Upper Water sample, and an Aquarium Solutions Accuprobe Hydrometer was used to estimate specific gravity and salinity of the sample.

### Limitations for Data

Methylene blue dye was chosen for its relatively low molecular weight (MW = 319.85 gmol^−1^). Due to its delocalized charge, it is a reasonable proxy for one product of CO_2_ absorption by MEA, the cation 2-hydroxyethylammonium, as both molecules electrostatically associate with the negatively charged carboxylate group on alginate. However, the dye is significantly more massive than MEA by a factor of 5.2, and therefore the rate of outward diffusion of this dye is a likely underestimation of the rate of outward diffusion for MEA.

## Results

After 6 additions of dry ice, varying in mass, 25 ml of calcium alginate beads containing 15% MEA absorbed 1.2 ± 0.1 grams of CO_2_. The theoretical maximum absorption of CO_2_ for the MEA present per trial (calculated from the 2:1 MEA:CO_2_ stoichiometric molar ratio), is 1.351 grams. This implies a high yield for CO_2_ absorption, suggesting little interference of the alginate hydrogel scaffold with the capacity of MEA for scrubbing carbon dioxide.

The absorptions of CO_2_ by mass for the MEA-alginate beads over all additions of dry ice were significantly higher than the blank containing 200 ml of distilled water. Final recorded masses of the solution containing the MEA-alginate beads and blank were compared to a secondary control containing a 200 ml solution of 1.5% CaCl_2_ (Fig. 3). As the beads were formed by ionic cross-linking of calcium, comparisons account for background absorption of CO_2_ by any free Ca^2+^ ions remaining in the solution (through formation of CaCO_3_ after H_2_CO_3_ formation). There were no significant differences between the secondary control containing CaCl_2_ and the blank containing distilled water, yet there were significant differences between the CaCl_2_ solution and the MEA-alginate beads, suggesting that the measured increase of mass is isolated to MEA-induced capture of CO_2_.

**Figure 2:**
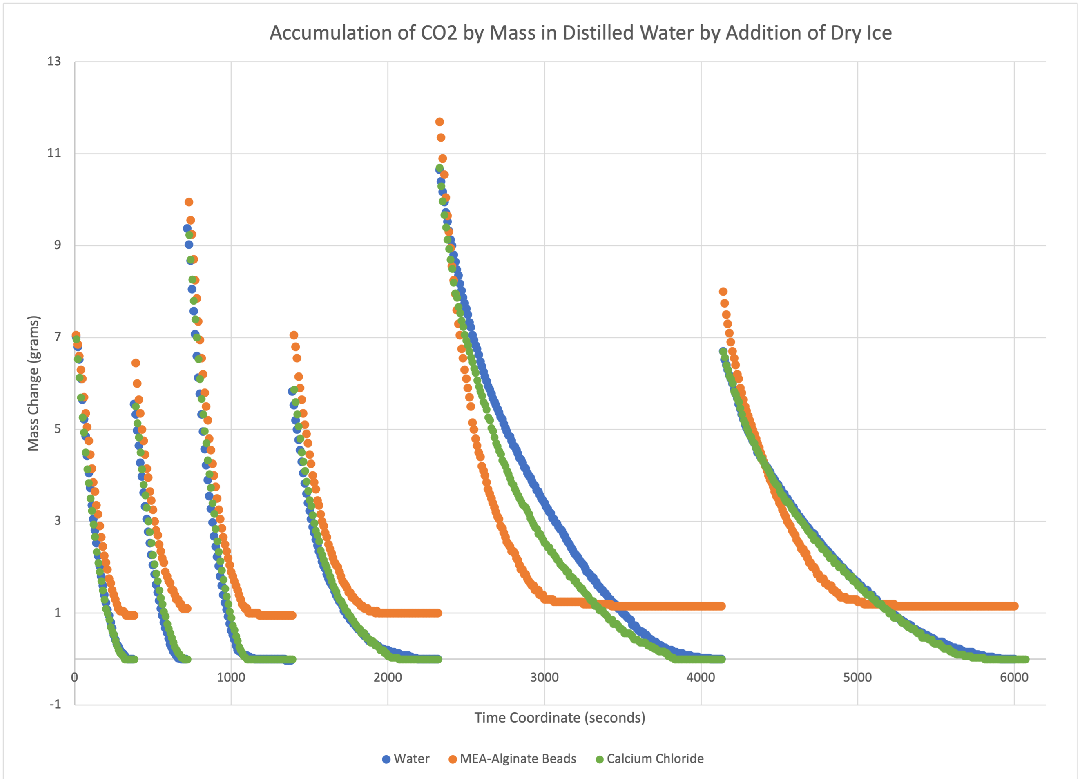
Mass Change recorded after 6 consecutive additions of dry ice to solutions containing 200 ml of water (blue), 1.5% calcium chloride (green), and a 200 mL mixture of water and 25 mL of MEA-encapsulated calcium alginate beads.

**Figure 3:**
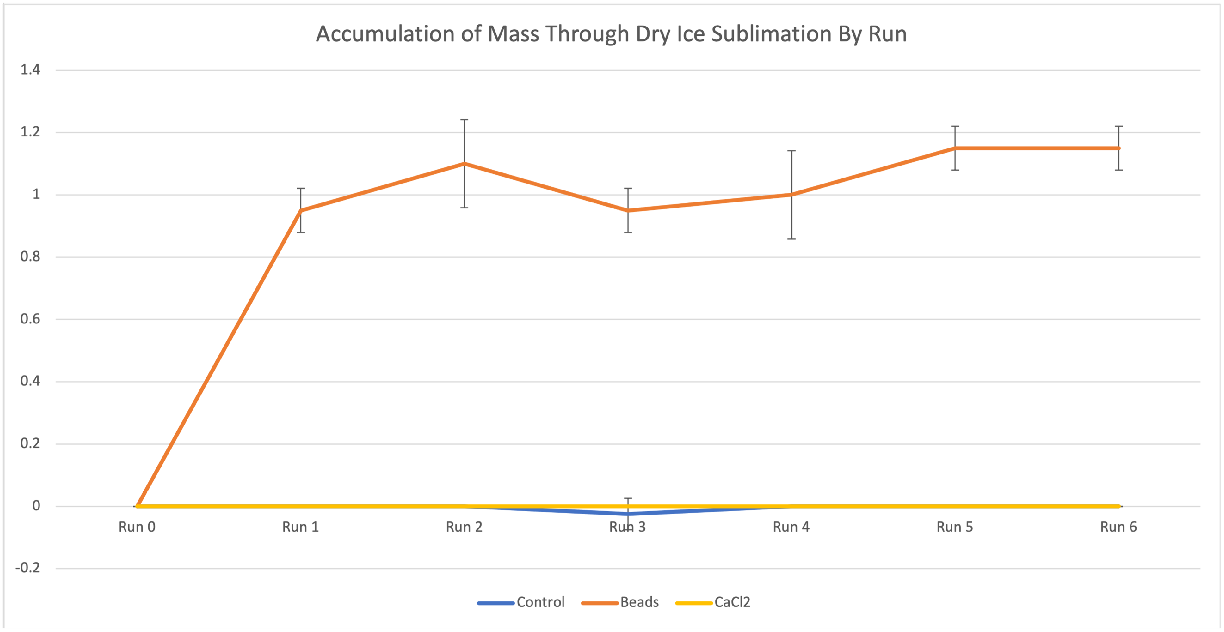
Net mass change recorded at the end of each run. Each run represents a subsequent addition of dry ice (the masses of dry ice added are recorded in supplementary information). Measurements for net mass change were taken 120 seconds after the dry ice had fully sublimated. Error bars represent standard deviation. Points with no error bars have a standard error smaller than mass balance uncertainty.

Sublimation time was estimated by measuring the total time taken for sublimation of each mass of dry ice, made audible through a microphone detecting sound conduction through the metal of the flask and mass balance as the dry ice sublimated. Trends in sublimation time per each dry ice addition remain constant yet diverge around halfway through the successive CO_2_ additions (Fig. 4). After this point, the trends between all conditions (MEA-alginate beads, CaCl_2_ solution, and blank conditions) diverge and remain diverged.

**Figure 4:**
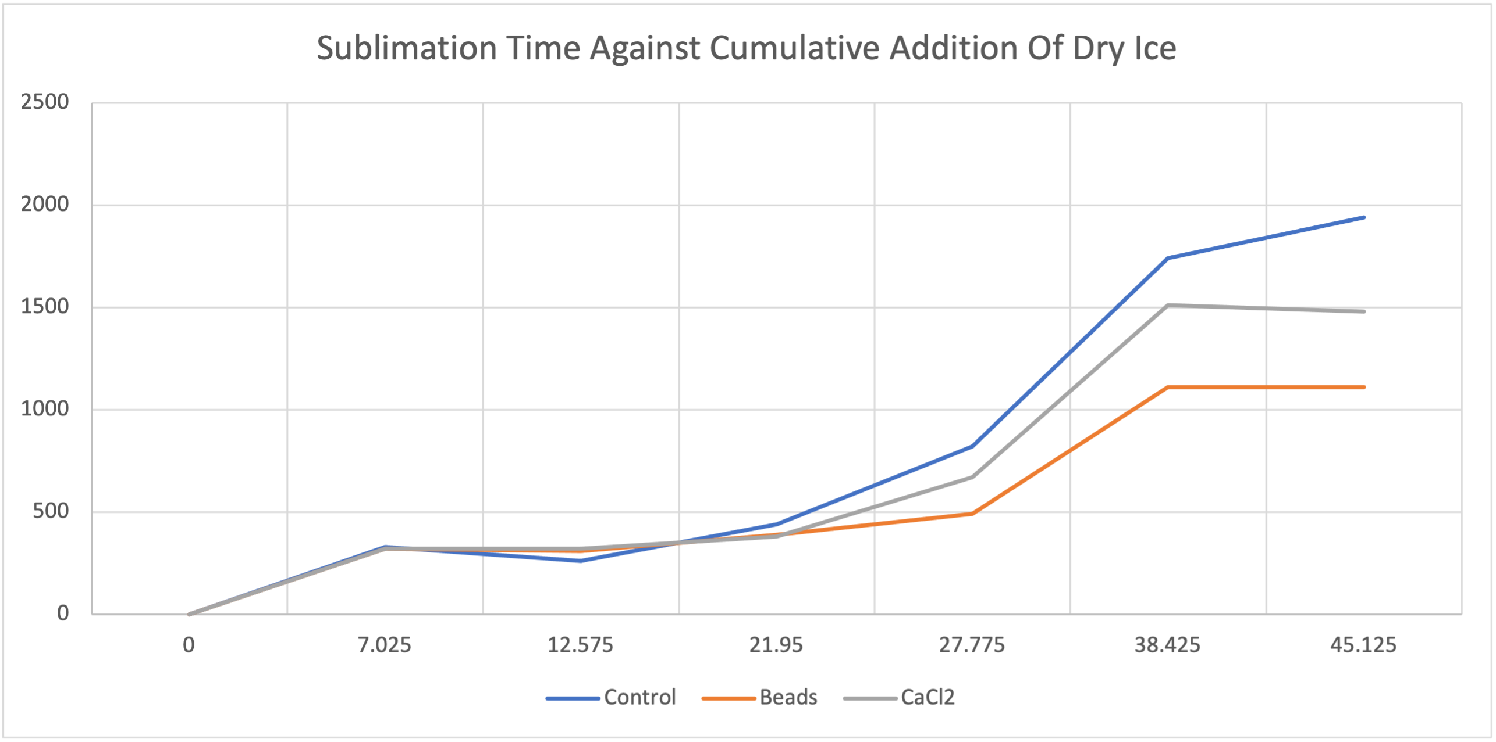
Sublimation time recorded against the cumulative mass of dry ice added for a blank control consisting of 200 ml of distilled water (blue), 200 ml of 1.5% calcium chloride (gray), and a 200 ml mixture of distilled water and 25 ml of MEA-encapsulated calcium alginate beads (orange). The x-axis represents cumulative mass of dry ice (grams), and the y-axis measures sublimation time (seconds).

Offering interpretation for the decreased sublimation time for the MEA-alginate beads in comparison to the CaCl_2_ and blank conditions; nonpolar CO_2_ is sparingly soluble in aqueous solution. CO_2_ is also electronically distinct from charged HOCH_2_CH_2_NH_3_^+^ or HOCH_2_CH_2_COO^−^, products of MEA-scrubbing of CO_2_ which are soluble in water as ionic compounds. Both the ethanolamine derivatives and CO_2_ have distinct, noninterfering solubilities in solution. MEA chemically sequesters CO_2_ into the soluble, charged forms, allowing more of the gas to dissolve at a faster rate through solution than a solution already saturated with CO_2_. Figure 4 therefore represents the “draining” of the carbon sink achieved by MEA (and, to a lesser degree, CaCl_2_) to allow for greater CO_2_ solubility in water.

The decreased sublimation time for the CaCl_2_ condition, relative to the blank condition, may be due to free Ca^2+^ ions increasing the CO_2_ absorption capacity of the solution through minor formation of calcium carbonate, while not able to form significant masses of the solid.

Figure 5 above measures the release of methylene blue dye through the core of the alginate beads through spectrophotometry. All beads sampled invariably displayed a net increase in absorbance, indicating that the dye ultimately leaked outwards through all conditions. The extent to which leakage occurred is suggested to be a product of both hydrogel concentration and layering.

**Figure 5:**
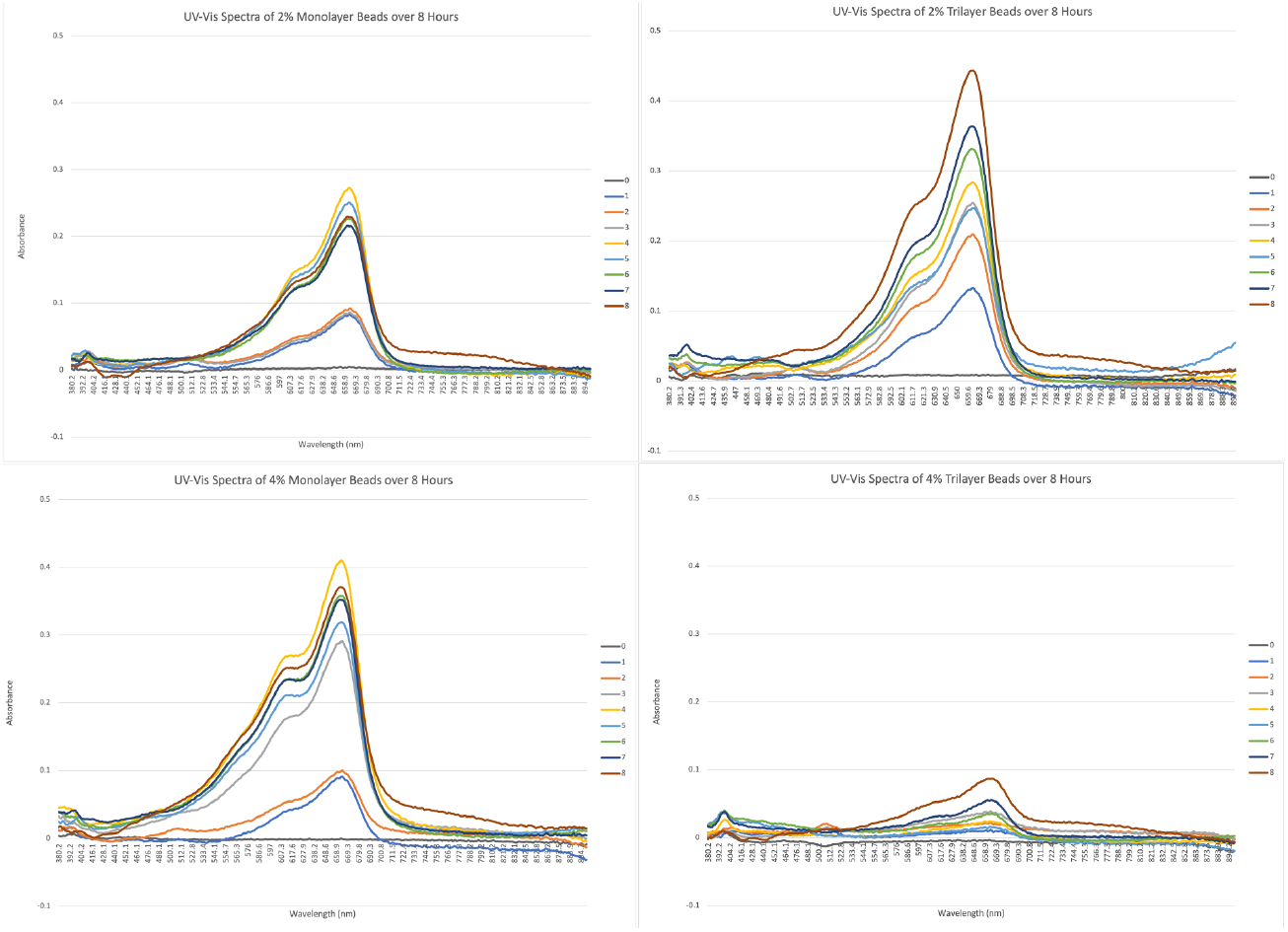
The matrix above displays overlayed hourly UV-Vis absorption spectra as methylene blue dye diffuses through the core of the alginate beads, over a period of 8 hours. The spectra shown are for a 2% alginate monolayer bead (top left), 2% alginate trilayer bead (top right), 4% alginate monolayer bead (bottom left), and 4% alginate trilayer bead (bottom right).

Prior research recognizes a burst-release profile for release through alginate hydrogel (Voo et al., 2016). Forming a monolayer alginate bead using a 2% alginate solution results in a burst release of dye, consistent with these findings. Interestingly, increasing the alginate concentration to 4% while still forming a monolayer bead displays a similar burst release profile. Maximum absorbance values for the 2% monolayer and 4% monolayer were both observed at 4 hours. When observations were made following the 8 hour period, two of three 2% monolayer beads and three of three 4% monolayer beads exhibited visible splitting.

Trilayering the alginate beads resulted in a release profile mimicking zero-order kinetics for constant, non-time-dependent release of dye. In contrast to the 2% and 4% monolayer beads, the maximum absorbance values for the 2% and 4% trilayer beads were both observed at 8 hours, indicating a relatively continuous release of dye through the hydrogel. The 4% trilayer beads displayed a significantly smaller magnitude of dye release in comparison to the 2% trilayer, 2% monolayer, and 4% monolayer beads. Decreasing the rate of release of substrate through the alginate hydrogel may be accomplished by balancing the increase of alginate concentration with increasing layering over the alginate-substrate core mixture.

The absorbance matrix above (Fig. 6) displays lower leakage (measured by absorbance at the λ_max_ of methylene blue) for beads prepared with ASUW (Atlantic Subarctic Upper Water) as a solvent beads placed in ASUW than when the same composition of beads were placed in distilled water. Additionally, beads prepared with distilled water as a solvent displayed lower leakage than when placed in ASUW. These results indicate that similar solvent composition of the beads to the aquatic environment they are placed in (namely, isotonicity) is necessary to reduce substrate leakage.

**Figure 6:**
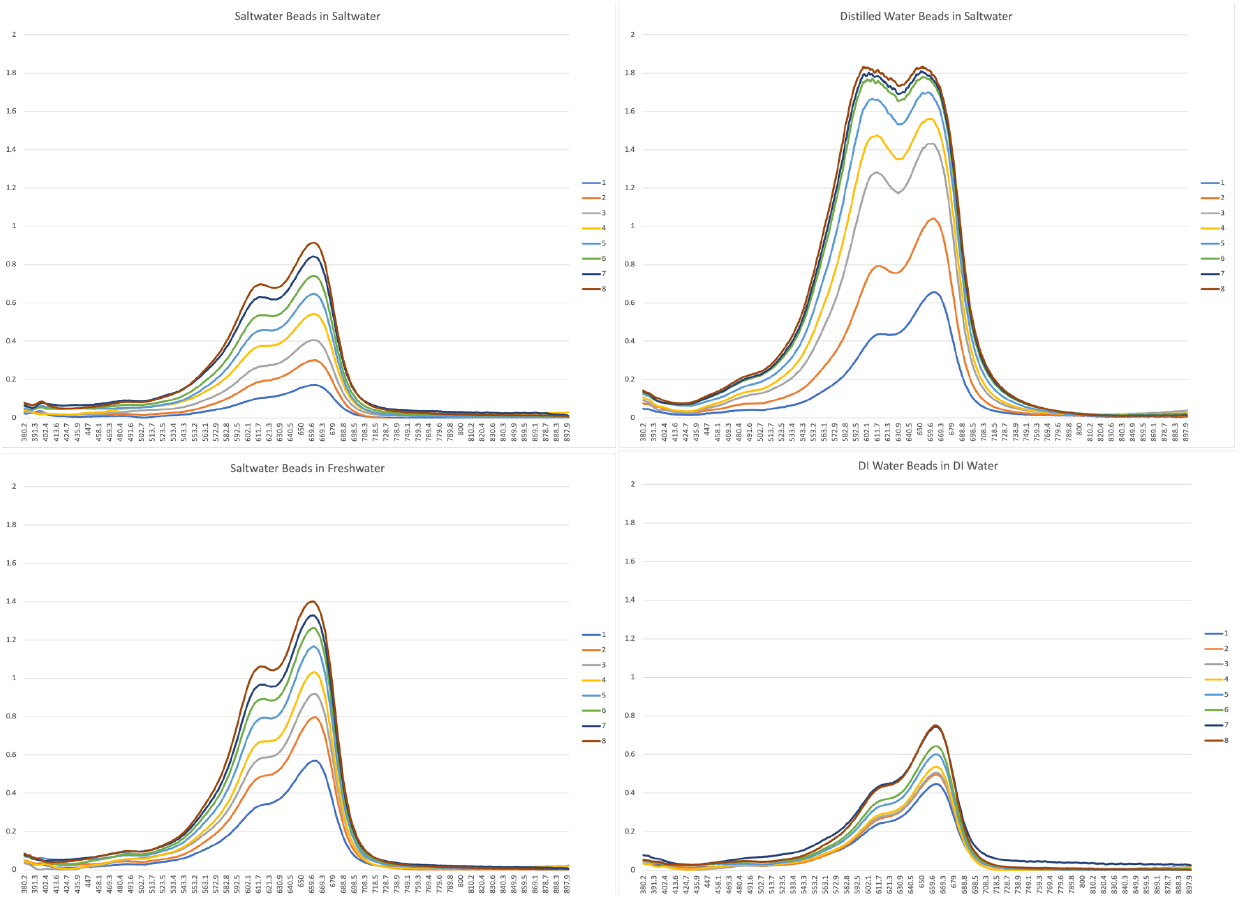
The matrix above displays hourly overlayed UV-Vis absorption spectra as methylene blue dye diffuses through the core of 2% trilayer alginate beads, measured over an 8 hour period. The beads in each condition differ by solvent used in preparation and solvent beads were placed in. The spectra shown above are for beads prepared in Atlantic seawater, placed in Atlantic seawater (top left), distilled water beads placed in Atlantic seawater (top right), Atlantic seawater beads placed in distilled water (bottom left), and distilled water beads placed in distilled water (bottom right).

Interestingly, while literature suggests the λ_max_ of methylene blue to be 667 nm, a blue-shift in the wavelength of maximal absorbance is witnessed in the presence of ASUW. The table above (Tab. 1) displays the λ_max_ of methylene blue after 24 hours in the corresponding aqueous phase identified above. The hypsochromic effects appears to be more pronounced in conditions with beads placed in solutions resulting in systems lacking isotonicity (i.e., blue-shift is more pronounced in ASUW beads in DI water than ASUW beads in ASUW). This may indicate potential side-reactions driven by increased dye concentration, as a product of outward diffusion of dyes due to the lack of isotonicity.

**Table 1:** Average λmax measured for alginate beads containing methylene blue dye under varying conditions. Beads were prepared in ASUW solvent and placed in ASUW or DI water, and beads were prepared with DI water solvent and dispensed in ASUW or DI water. Corresponding deviation from literature λmax for methylene blue, 667 nm, is indicated in the bottom row, indicating the magnitude of blue-shift in each condition. Blue-shift is the most pronounced in columns 2 and 3, where gel beads were placed in solutions hypertonic or hypotonic, respectively, to the gels.

| Condition / $\lambda_{\max}$ values | ASUW beads in ASUW | DI Water beads in ASUW | ASUW beads in DI Water | DI Beads in DI Water |
| --- | --- | --- | --- | --- |
| $\lambda_{\max}$ after 24 hrs in solution (nm) | 660.4 | 651.5 | 653 | 662.6 |
| $\Delta\lambda_{\max}$ from literature (nm) | -6.6 | -15.5 | -14 | -4.4 |

As blue-shift occurs in all solutions, regardless of DI or ASUW solvent, the increased availability of dye in a surrounding aqueous phase may have increased the conversion of the dye into alternative forms with smaller wavelengths for maximal absorbance. One such alternative form is the dimerized form of methylene blue, abbreviated (MB^+^)_2_ which exhibits a blue-shifted λ_max_ of 608 nm in comparison to 667 nm for the monomer (Morgounova et al., 2013).

Osmotic drive of the dye out of the beads in hypotonic solutions would have increased the apparent concentration of dye, offering more availability of monomer to dimerize; this may account for the trend of increased blue-shifting in anisotonic systems.

### Negative Findings

The degree of leakage of MEA from alginate beads was realized in an exploratory experiment, after forming 25 ml of 2% calcium alginate beads in 1.5% CaCl_2_ solution. This mixture was stirred using a magnetic stirrer at high speed for 10 minutes. The alginate beads were macroscopically intact after this period, and the surrounding solution was transferred to a vacuum-sealed insulated flask. Accumulation of CO_2_, measured by consecutive additions of dry ice, was measured to be significantly higher than the blank, indicating a significant degree of leakage from the beads after this 10-minute agitation. This observation revealed that leakage — due to the small molecular weight of MEA — would restrict MEA-encapsulated calcium alginate beads to carbon capture and storage in the short term, due to eventual outward diffusion of the small molecule.

This prompted the investigation into multilayering to prolong the effectiveness of these beads by controlling the rate of outward diffusion.

## Discussion

### Biosynthesis for Monoethanolamine

Monoethanolamine is a derivative of the non-essential amino acid serine (2-amino-3-hydroxypropanoic acid), being the decarboxylated form of the amino acid. While the amine is industrially synthesized by reacting ammonia with ethylene gas, MEA may also be biologically synthesized using an enzyme that catalyzes the decarboxylation of serine. Serine decarboxylase, or SDC is an enzyme found in *Arabidopsis thaliana* (Mouse-ear cress). SDC works specifically on the L-serine enantiomer to decarboxylate it, forming monoethanolamine and carbon dioxide as a byproduct.

SDC does not participate in feedback inhibition with its products, CO_2_ or HOCH_2_CH_2_NH_2_. As monoethanolamine or carbon dioxide does not inhibit the action of SDC, the presence of L-serine will elicit the complete specific decarboxylation of this substrate.

This method encouragingly differs from the industrial method to produce ethanolamines in that, provided the enzyme SDC is not exposed to denaturing conditions, a sample of the enzyme can be reused repeatedly. As the enzyme is chemically unaffected and cannot be inhibited by the product, the catabolizing process is also efficient. Due to L-serine being non-essential, the amino acid is produced by numerous organisms. Encouraging the expression of genes which synthesize enzymes related to serine production (such as *E. Coli* genes serA, serB, and serC) would promote the hyperproduction of L-serine, a suitable precursor for the synthesis of MEA. Mundhada et al. produced high-yield E. coli strains by knocking out serine deaminases and a transferase; deletion of genes prevented the degradation of serine to glycine and pyruvate, which allowed for anaerobic serine production by fermentation at 0.43 grams serine per gram of glucose, the highest reported yield to date (Mundhada et al., 2016). The buildup of serine toxicity reported in the literature can be mitigated by mild SDC treatment to the growth medium, which may allow for direct extraction of MEA from the source.

### Microalgae as a CO_2_ Scrubbing Agent

One clear benefit to carbon dioxide scrubbing through immobilization of the scrubbing agent in alginate hydrogel scaffolding is that a variety of scrubbing agents can be used. Monoethanolamine was chosen for its wealth of publicized data, mainstream use in industry, and biosynthetic potential. However, due to its low molecular weight, it can easily pass through the pores of calcium alginate and diffuse through a single layer of calcium alginate rapidly. This problem was mediated in the short-term by utilizing repeat gelation to assemble a bead with three distinct alginate layers, which reduced the rate of dye dissolution only when paired with an increased alginate concentration (Fig. 5).

However, scrubbing agents larger than the alginate pore size, such as photosynthetic microalgal species or cyanobacteria, may appear to close the gap. Alginate beads consisting of Gram-positive bacteria *Bacillus subtilis* have been prepared in the past to treat wastewater sewage (Su et al., 2021). Given the relatively larger size of macroalgal species such as chlorella and spirulina, long-term entrapment by biological sequestration may be possible. Further research and clarification on the sustainability of such a bead may prove necessary.

### Afforestation for Macroalgae Forests

As CO_2_ capture using immobilized MEA has been demonstrated, it is important to consider the factors necessary for financial viability.

Primarily, symbiotic relationships between algae and bacteria have been evaluated in the past and would serve an effective measure for the integration and amplification of the proposed method on a global scale (Higgins & VanderGheynst, 2014; Ramanan et al., 2016). By building an algae and modified E. coli photobioreactor (PBR), a mutualistic interdependence between the serine-producing E. coli and the macroalgal species can decrease the strain on extraneous resources, such as bacterial growth medium or algal fertilizer.

Brown algae (Phaeophyceae) is a class of photosynthetic algae notable for its high rate of growth and the composition of cell walls, which contain a high concentration of alginate. One proposed genus is *Macrocystis*, a large, monospecific genus of algae. The species *Macrocystis pyrifera* is one of the primary sources of alginate or the salts of alginic acid, and therefore a source of great interest for this paper.

By growing the algae in a PBR with the modified E. coli (Fig. 7), the two species can symbiotically coexist with significantly less external regulation than that required for cultivating the species alone. Alginate from the algal cell wall and MEA produced by the modified, SDC-treated bacteria can both be extracted. Excess oxygen produced by brown algae photosynthesis is released as a by-product into the atmosphere.

**Figure 7:**
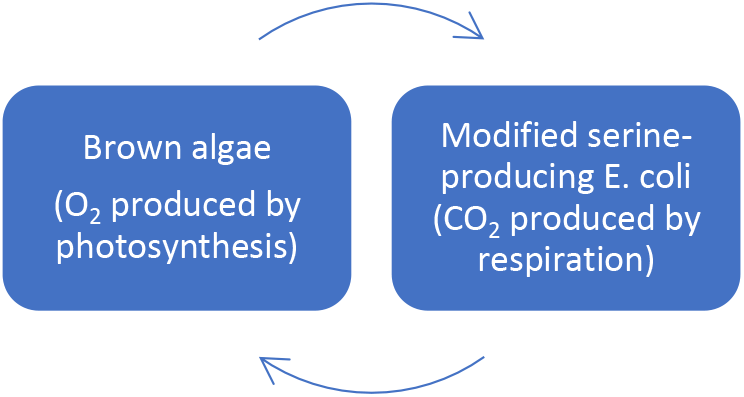
Mutualistic interdependence through vital gas exchange between algal species producing alginate and modified E. coli species producing L-serine, harvested to generate more MEA.

*Macrocystis* also grow well in cold aquatic climates. This may allow for proximity of the PBRs to locations of ideal carbon scrubbing temperatures, reducing the financial cost of hydrogel transportation.

In addition to the added capacity for carbon capture, afforestation using *Macrocystis*, supported by CO_2_ capture-functionalized alginate hydrogel can rapidly restore previously degraded kelp forests, stabilize aquatic trophic levels by providing shelter for primary consumers and maintain the aquatic ecosystem’s biodiversity. This outcome cannot only be accomplished by *Macrocystis* but also by algal species noted in Fig. 8 (*Nereocystis*, *Lessonia*, *Laminaria*, and *Ecklonia*). As many coastal regions have kelp forests in proximity, these sources for on-site sequestration are readily available, should integration on a global scale be pursued.

**Figure 8:**
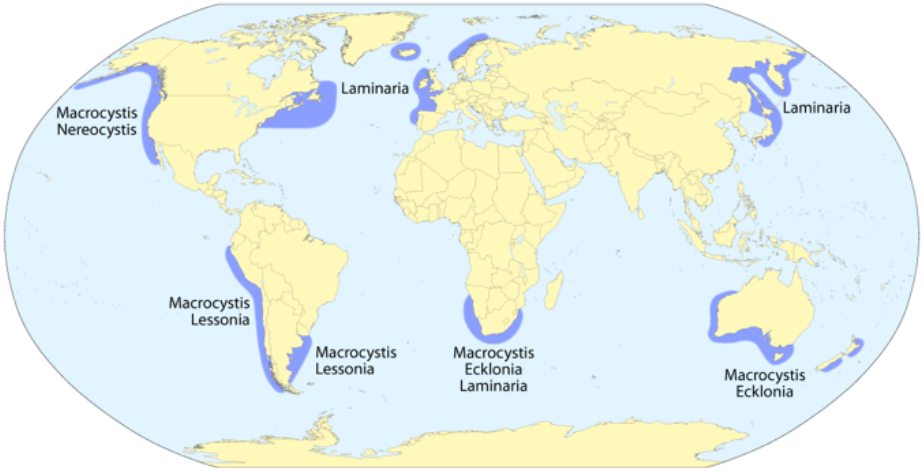
Locations and dominant genera of the Earth’s distributed kelp forests (Dörrbecker, 2011).

Additionally, the locations of kelp forests (Fig. 8, Dörrbecker, 2011) provide substantial information for where to produce and deploy immobilized MEA. Polar coastal regions (eastern and western coasts of North America, the southern tip of Africa, the southern borders of Australia and New Zealand, and the eastern and western coasts of Eurasia) are locations of great kelp forest density. As kelp grow in cold climates (Steneck et al., 2002), many of these locations could be ideal for carbon capture and eventual biological sequestration within the algae.

While the integration of brown algae afforestation and L-serine production reduces the strain for additional cultivation resources, there are still some factors necessary for the process of immobilization. For instance, a source of calcium is necessary to form the insoluble calcium alginate, to immobilize MEA. As alginate is documented by literature to preferentially uptake large-radius heavy metal ions such as Pb^2+^ from the aqueous phase in exchange for Ca^2+^ release into solution, calcium can therefore be recycled back into a production line for further gel production, while entrapping divalent heavy metals within a gel as a sequestered waste product (Wang et al., 2016). Ion-ion exchange may present a means for recycling calcium, a relatively safe divalent metal, into new alginate gels, prompting greater sustainability.

### Aquatic Carbonate Minerals and Ocean Acidification

The oceans and most other bodies of water are undergoing acidification. Acidification is one consequence of increased concentrations of dissolved CO_2_ in the ocean. As dissolved CO_2_ forms carbonic acid, the increase in atmospheric CO_2_ levels directly correlates with the decrease in aquatic pH. This increase in dissolved CO_2_ concentrations, therefore, correlates directly with the decrease in aquatic concentrations of aragonite — a key product of the calcification process that aquatic shell-building organisms undergo.

Aragonite, calcite and vaterite are carbonate structures that aquatic organisms can use to biogenically build their shells. As the ocean acidifies, protons from introduced carbonic acid degrade these carbonate structures, releasing more carbonic acid into the ocean.

By capturing carbonic acid in the form of CO_2_, the acidic effect of increased carbonic acid can be reduced, promoting calcification. This would therefore allow for atmospheric CO_2_ to dissolve in the oceans, which can be captured in the same manner. Carbonate structures such as aragonite also play a role in scrubbing pollutive substances such as lead, which have the potential to accumulate and destabilize aquatic ecosystems (Köhler et al., 2007).

### Synthetic Seeds and Biological Sequestration

Outside of the aquatic ecosystem, applications of alginate beads in agriculture have been extensively explored. Synthetic seeds can be formed by implanting plant embryos or complete seeds into alginate beads, providing a hydrating environment with which embryos may grow and seeds may germinate. The transparency of the hydrogel provides a source of light for photosynthetic reactions, while the high specific heat of water, within the hydrogel, provides a degree of thermal resistance.

Synthetic seeds have been formed extensively in past literature, and the viability of embryos while producing crops such as sweet corn (Thobunluepop et al., 2009), potatoes (Nyende et al., 2002) and rice (Kumar et al., 2005) from synthetic seeds have been evaluated. In locations where shortages for food production are prevalent, synthetic seeds may be worthwhile for not only growth, but also mass transport of seeds, as the alginate-based synthetic seeds may lengthen embryo viability.

Ammonia has been evaluated recently as a carbon dioxide scrubber which offers key advantages over MEA. Ammonia is a synthetic precursor to MEA, and is produced in mass through reactions such as the Haber-Bosch process (Luis, 2016). In the scrubbing reaction with CO_2_, the product is ammonium carbamate, which exists in equilibrium with urea – a key fertilizer for plant growth – with water. Using ammonia as a CO_2_ scrubbing agent within alginate beads offers the potential for urea to provide a fertilizing environment for synthetic seeds, providing a means for direct biological sequestration of absorbed CO_2_ by plants. While substrate leakage is indeed a recurrent problem for ammonia as it is for MEA, the use of these beads for short-term capture and biosequestration is possible — presenting implications for aiding not only terrestrial afforestation but aquatic afforestation.

## Conclusion

In this paper, we have presented a novel method for biocompatible carbon dioxide capture in aquatic ecosystems.

Coupling afforestation techniques, using rapidly growing species of brown algae, with the potential for symbiotic biosynthesis of monoethanolamine by cellular anaerobes may present a financially viable means for large-scale application of such a method. This method has the potential to boost the CO_2_ absorption abilities of these ecosystems much more significantly than using aquatic or terrestrial afforestation alone.

Particularly significant work has been conducted recently concerning the effectiveness of amine scrubbers such as MEA and the specific capturing ability of carbon dioxide by these scrubbers. This paper utilizes the foundation of this knowledge to ultimately detail the development of a functionalized, biocompatible hydrogel which can capture carbon dioxide with a relevant degree of safety. As proposed in the paper, the method is functional, effective, and scalable.

Prior literature recognizes the drive for reorienting carbon capture modes toward developing closed processes wherein CO_2_ products have economic value and materials can be recycled or regenerated for continued use (Luis, 2016). As such, a switch to from MEA to ammonia may present as an opportunity to develop fertilized synthetic seeds following CO_2_ capture.

This work is significant not only in the context of counteracting the effects of global warming but also in terms of determining a means for supporting and stabilizing the carbon cycle in the future. It is undoubtable that even while a source of fully renewable energy may be made the primary source of energy, carbon will need to be captured indefinitely. This method may prove worthy of carbon capture through its use in an aquatic environment and may retain the capacity to significantly drain carbon sinks if upscaled.

This is not to suggest that the proposed method is without flaws. The cost of transportation of the immobilized product to and from aquatic regions is a significant obstacle to overcome, and research in deployment and recovery is necessary and encouraged. The choice of scrubbing agent largely dictates the length of its effectiveness, and benefits must be weighed against accessibility and affordability of these materials. The construction of such photobioreactors in the locations suggested previously in Figure 6 will require major international funding and support.

However, with considerable investment in the application of this method, continued research into further means of carbon dioxide recyclability, integration into our current means for production, and global, multipartisan engagement, the largely devastating consequences to the environment, which we are already witnessing, can be counteracted.

## Acknowledgements

The author would like to thank the Laidlaw Foundation for funding this project. The author would also like to thank Professor Ged Parkin of Columbia University, and all members of the Parkin group for their continued support and assistance in carrying out the experiments.

## Supplementary Materials

### Supplementary Graphs (Enlarged)

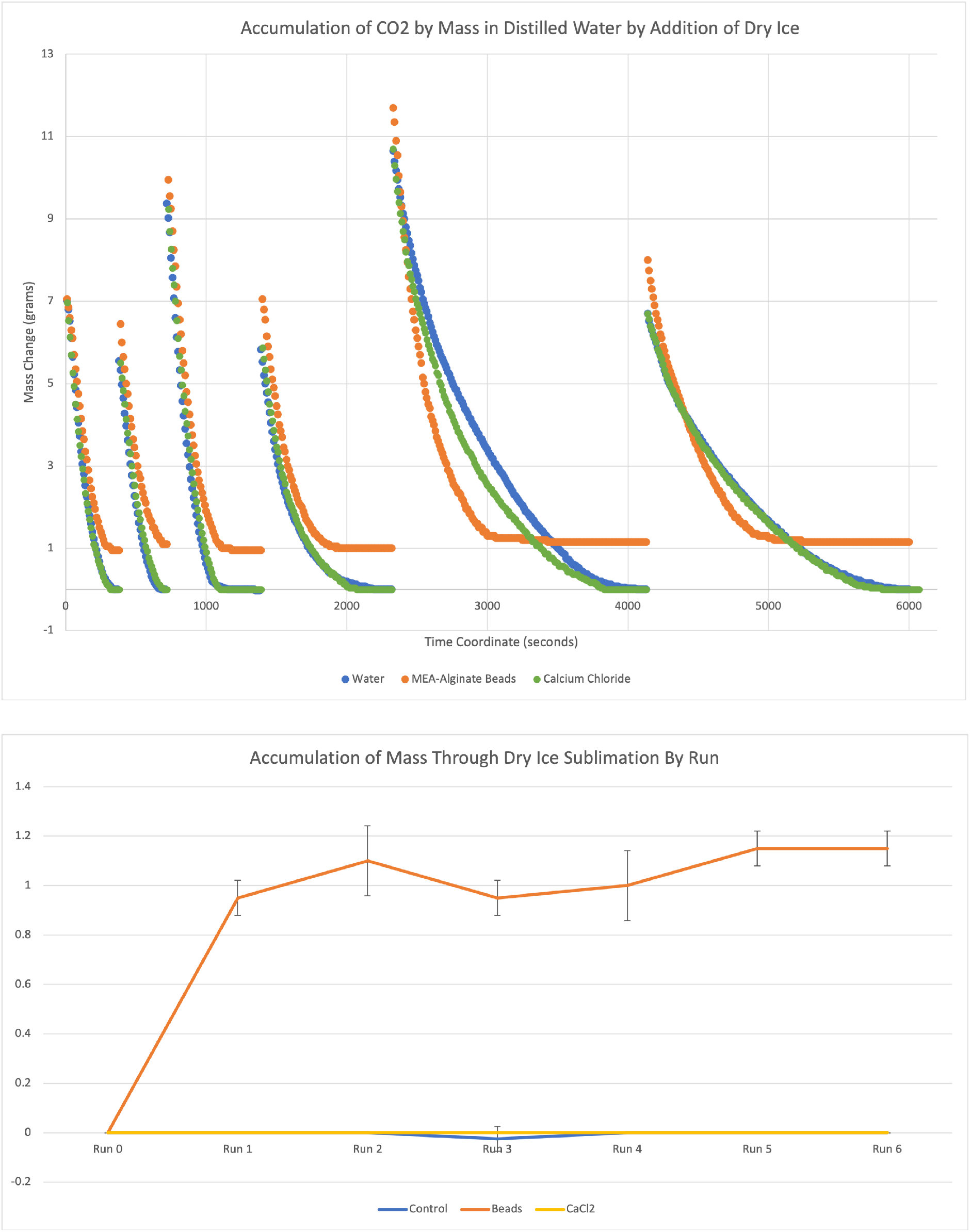

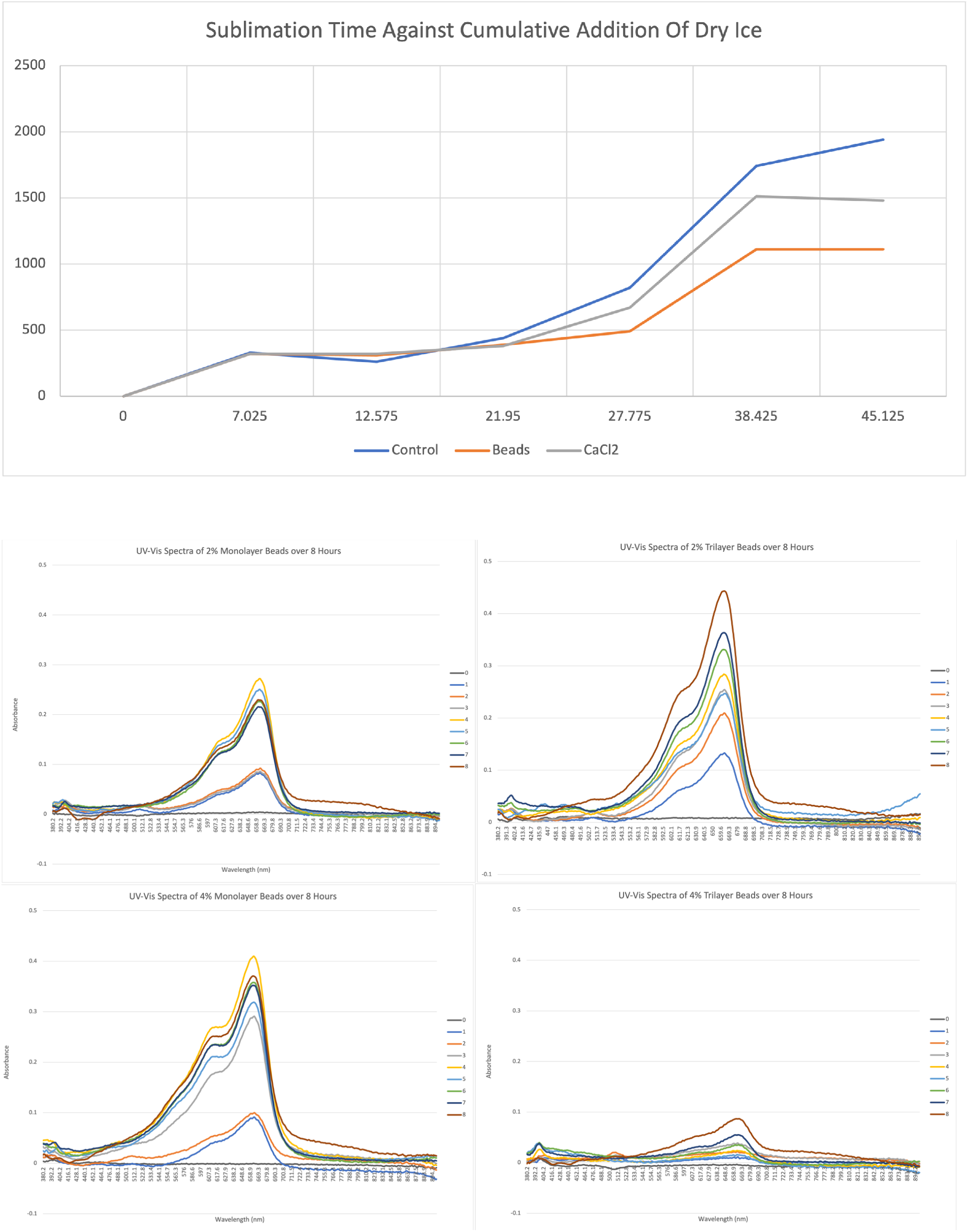

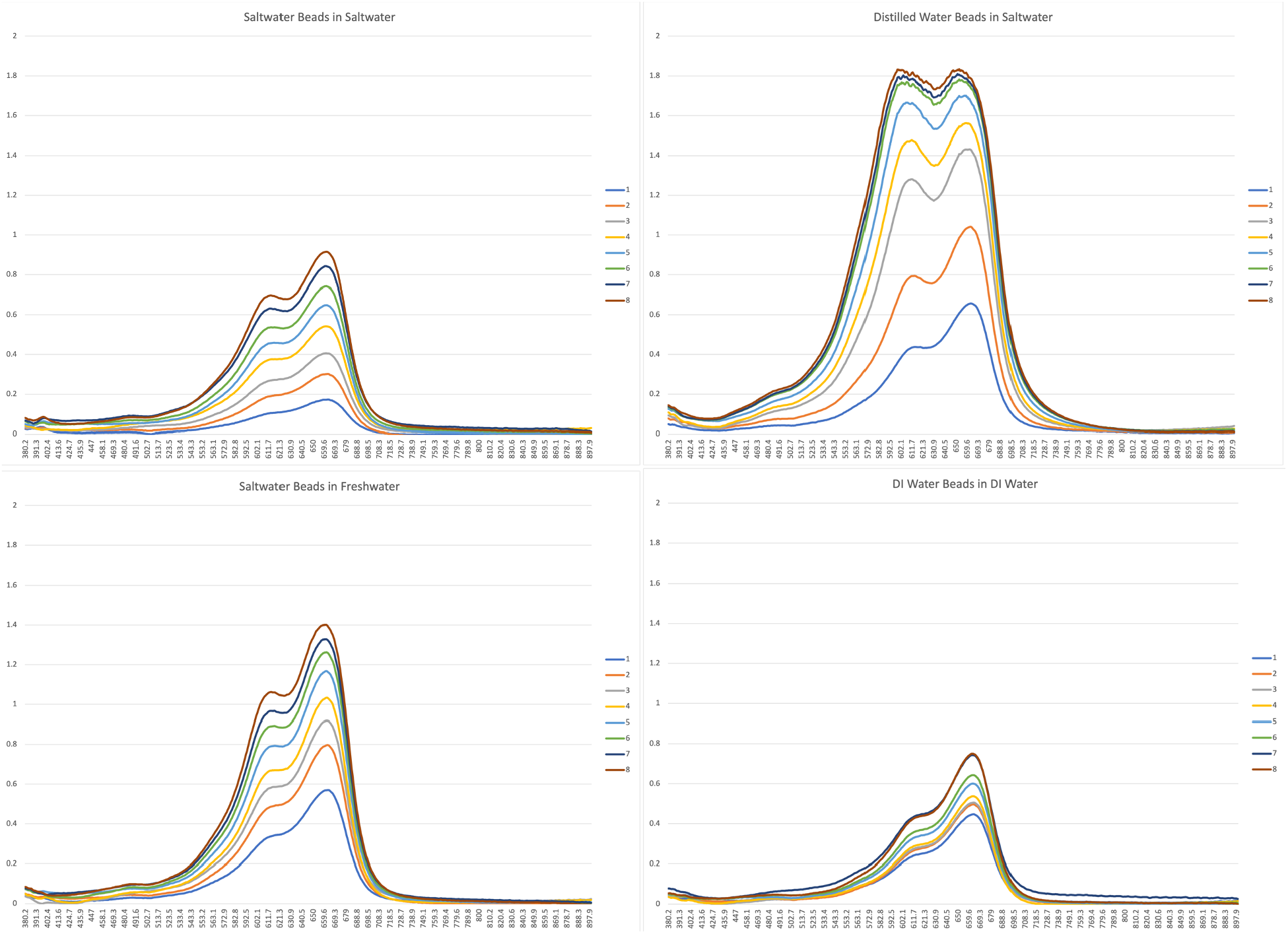

### Supplementary Table

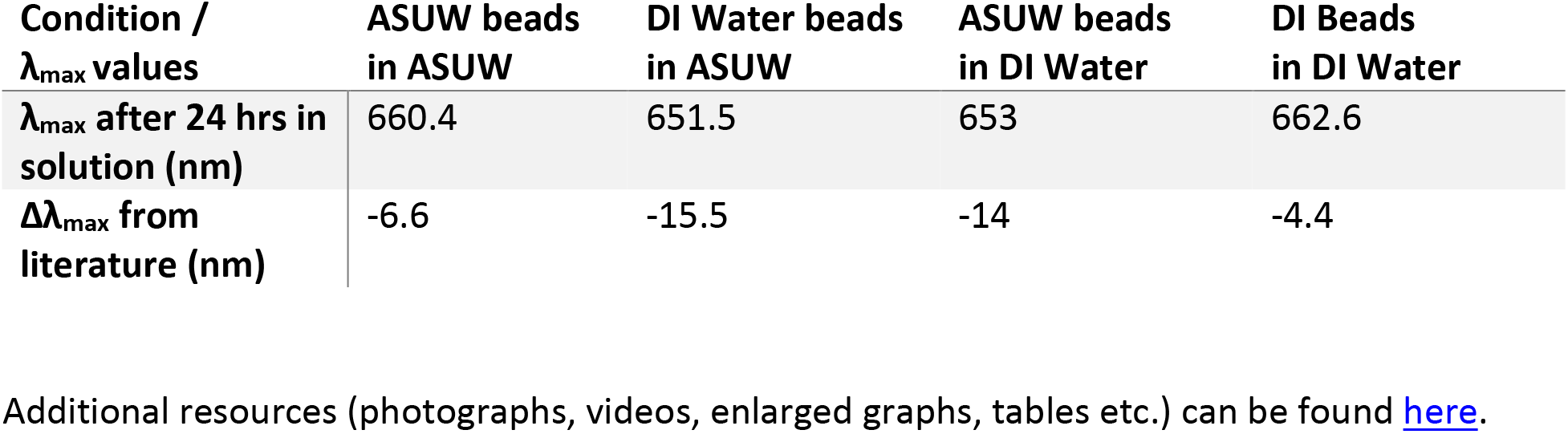

| Condition / $\lambda_{\max}$ values | ASUW beads in ASUW | DI Water beads in ASUW | ASUW beads in DI Water | DI Beads in DI Water |
| --- | --- | --- | --- | --- |
| $\lambda_{\max}$ after 24 hrs in solution (nm) | 660.4 | 651.5 | 653 | 662.6 |
| $\Delta\lambda_{\max}$ from literature (nm) | -6.6 | -15.5 | -14 | -4.4 |

